# Functional localization and categorization of intentional decisions in humans: a meta-analysis of brain imaging studies

**DOI:** 10.1101/2020.11.27.401208

**Authors:** Ruoguang Si, James B Rowe, Jiaxiang Zhang

**Affiliations:** Cardiff University Brain Research Imaging Centre, School of Psychology, Cardiff University, Cardiff CF24 4HQ, United Kingdom; Department of Clinical Neurosciences, University of Cambridge, Cambridge CB2 0QQ, United Kingdom; Medical Research Council Cognition and Brain Sciences Unit, University of Cambridge CB2 7EF, United Kingdom

**Keywords:** intentional decision, free choice, meta-analysis, ALE, fMRI, PET

## Abstract

Brain-imaging research on intentional decision-making often employs a “free-choice” paradigm, in which participants choose among options with identical values or outcomes. Although the medial prefrontal cortex has commonly been associated with choices, there is no consensus on the wider network that underlies diverse intentional decisions and behaviours. Our systematic literature search identified 39 fMRI/PET experiments using various free-choice paradigms, with appropriate control conditions using external instructions. An Activation-Likelihood-Estimate (ALE) meta-analysis showed that, compared with external instructions, intentional decisions consistently activate the medial and dorsolateral prefrontal cortex, the right insula and the inferior parietal lobule. We then categorized the studies into four different types according to their experimental designs: reactive motor intention, perceptual intention, inhibitory intention and cognitive intention. We conducted conjunction and contrast meta-analyses to identify consistent and selective brain activations within each specific category of intentional decision. Finally, we used meta-analytic decoding to probe cognitive processes underlying free choices. Our findings suggest that the neurocognitive process underlying intentional decision incorporates anatomically separated components subserving distinct cognitive and computational roles.

## 1. Introduction

To fulfil our goals or desires, we constantly interact with the external environment through our voluntary behaviour. In contrast to reflexes that are beyond volition (e.g., a knee-jerk reflex), voluntary behaviours are characterised by choice (Passingham, 1995). Volition characterises the intentional choice or decision between multiple options, where the choice is not sufficiently explained by differences in expected or explicit rewards. The concept of intentional decision refers to this fundamental ability of human cognition: acting voluntarily based on internal or endogenous intentions (Marken, 1982).

The role of intention in decision-making occupies a broad spectrum. At one extreme lies externally guided perceptual decision such as stopping at a red traffic light, for which the involvement of internal intention is low because learned rules can dictate a correct choice (even if one can voluntarily break such rules). At the other extreme lies improvisational behaviour in music, painting or dance, which can be strongly determined by moment-to-moment intention. In between lies the common scenario of intentional decision-making, where the external environment constrains only which options are available while internal intentions dictate which of those options to choose. The ability to choose actions, cognitive strategies and behaviours in this way plays a key role throughout the life span and is essential to our understanding of human cognition. In child development from birth to 12 months, actions such as grasping and its coordination with vision gradually emerge from simple reflexes (Piaget, 1976; Beilin and Fireman, 1999; Lewis, 2010). In patients with neurodegenerative disorders, the inability to engage appropriate intentional behaviour can manifest as apathy (Starkstein et al., 2001), impulsivity (Dalley et al., 2011) and perseveration (Hughes et al., 2013). In addition, intentional behaviour is a foundation of social interactions via cooperation and collaboration (Bratman, 2017).

Intentional actions have been characterised by three components in the *what*-*when*-*whether* (WWW) model: (1) *what* action to perform, (2) *when* to perform it, and (3) *whether* to perform the chosen act (Brass and Haggard, 2008). The WWW model is based on evidence from two interlinked lines of research. First, the *when* component has been investigated by examining neural signatures immediately prior to intentional actions. Libet’s intentional action paradigm is a classic example of this type (Libet et al., 1983; Libet, 1985), which has been used to localize electrophysiological and BOLD activity in the medial-frontal cortex preceding the conscious awareness of subsequent voluntary actions (Lau et al., 2004a; Fried et al., 2011) (but see Trevena and Miller, 2002; Nachev and Hacker, 2014 for critical evaluations). Second, research on the *what* and *whether* components, the focus of the current study, commonly use variants of the “free-choice” paradigm^1^, to determine the neurocognitive mechanisms of voluntary decision processes.

In a typical free-choice paradigm, participants make a voluntary choice from multiple alternatives on each trial. The available alternatives can either be similar to each other (e.g., responding with different fingers, Zhang et al., 2012) or distinct (e.g., to choose voluntarily between stopping and acting in the adapted Go/NoGo task, Karch et al., 2009). Importantly, participants are made aware that all available options are homogeneous interms of their objective outcomes, and the tasks do not introduce or manipulate rewards of costs according to the choices made. In other words, the task is not to identify a correct response. Rather participants can choose any of the available options. The alternate options are equally appropriate, and one’s decision must come from intention. The intention could be influenced by endogenous factors, including subtly differential effort, preferences (Zajkowski et al., 2020), habits (Graybiel, 2008), incorrectly inferred arbitrary rules for the task, and recent actions (Zhang and Rowe, 2015; Phillips et al., 2018).

In recent years, there has been a substantial number of brain imaging studies adopting free-choice paradigms, enabling a well-powered meta-analysis. The current study focused on the hemodynamic and metabolic contrasts of intentional choice vs. specified response, which is the most widely reported task-related effect across free-choice studies. Here, specified responses serve as a control condition, in which participants need to make specific responses determined by the experimenter, rather than choose voluntarily from the same set of options in the free-choice condition. Therefore, the contrast between the two conditions offers an imaging maker of brain activation associated with intentional behaviour, controlling for the common effects of stimulus encoding and response initiation.

The objectives of this study were three-fold. First, to identify brain regions consistently activated by intentional decision, we performed a systematic search of BOLD-fMRI or PET studies of intentional decision and conducted an activation likelihood estimation (ALE) meta-analysis. Increased BOLD and PET responses during intentional choices are commonly reported in a frontoparietal network centred on the medial frontal cortex (Brass and Haggard, 2008). However, some studies also observed activations external to this network during intentional behaviour, in particular in the insula (Brass and Haggard, 2010; Thimm et al., 2012; Dall’Acqua et al., 2018) and the inferior frontal gyrus (Wisniewski et al., 2016). Because results from a coordinate-based ALE meta-analysis are pooled from a large number of participants in multiple studies, they usually have higher statistical power than a single experimental study (Walker et al., 2008).

Second, we conducted further contrast and conjunction meta-analyses, assessing the distinct and overlapping neural correlates between different types of intentional behaviour. As highlighted above, the nature of options in a free-choice paradigm can vary significantly between studies and hence involve different cognitive processes. We reviewed all studies to date that met our predefined inclusion criteria (see *Study selection and inclusion criteria*). Based on the experimental design and implementational details of individual studies, we proposed four categories of the free-choice paradigm (Figure 1): reactional intention (RI), perceptual intention (PI), inhibitory intention (II) and cognitive intention (CI).

**Figure 1.**
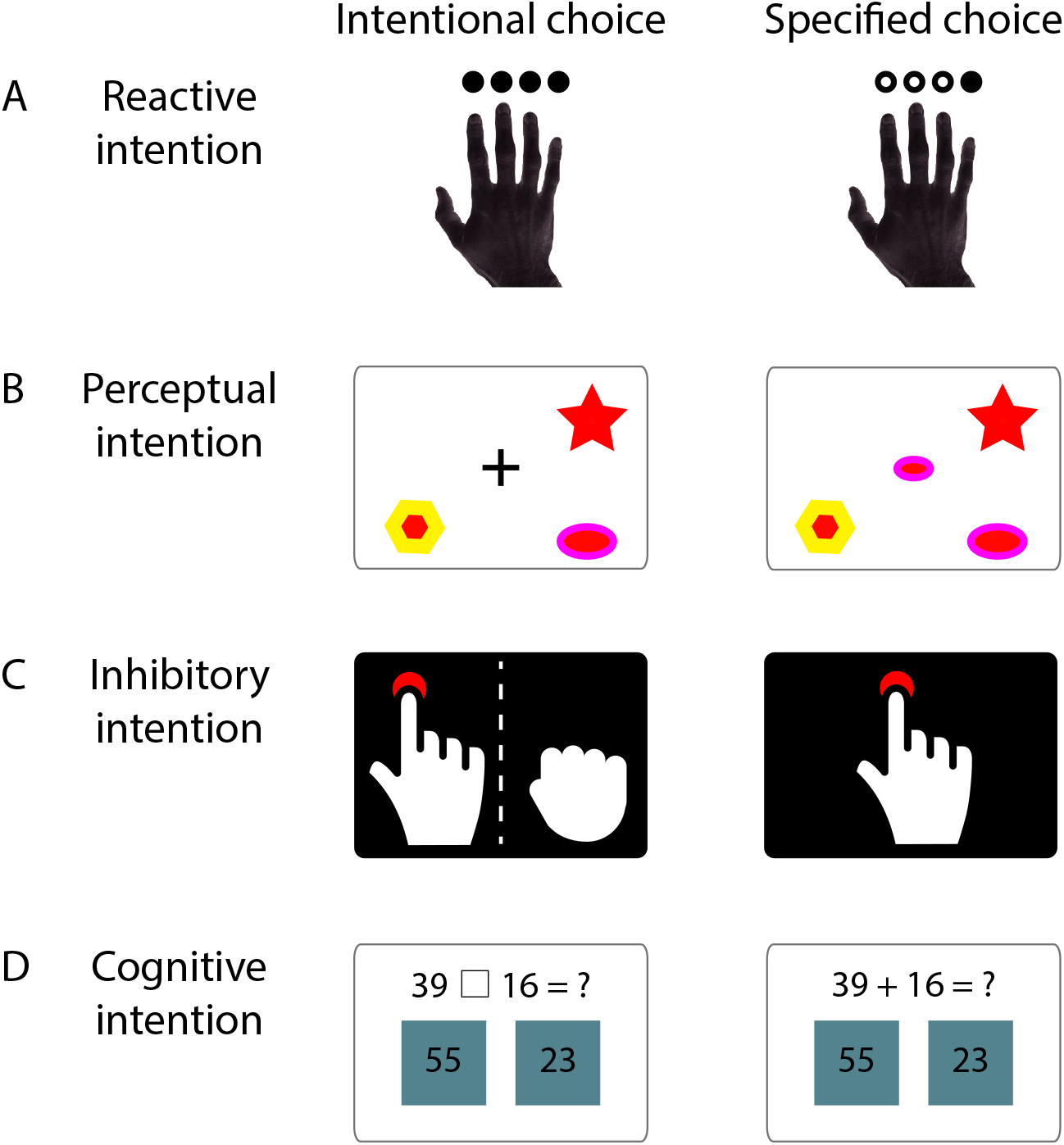
Schematics of four categories of free-choice studies. (A) In the reactive intention (RI) paradigm, task cues indicate directly available options (e.g., Rowe et al., 2008). (B) In the perceptual intention (PI) paradigm, task cues contain perceptually similar options that associated with different options (e.g., Lau et al., 2004). (C). In the inhibitory intention (II) paradigm, one of the options is to abandon or abort an intended action, and hence participants make voluntary choices between Go and Stop (e.g., Dall’Acqua et al., 2018). (D). In the cognitive intention (CI) paradigm, participants choose between different operations that require higher-level cognitive processing (e.g., Taylor et al., 2008). Behavioural responses are dependent on the execution of the chosen operation.

Third, to undertake an exploratory data-driven analysis, testing whether consistent BOLD-fMRI/PET patterns of intentional behaviour correspond to specific cognitive processes. We quantified the similarity between the meta-analytical whole-brain activation pattern estimated from free-choice studies and brain activation patterns from 100 specific cognitive topics, extracted from a database of over 11,000 brain imaging studies (Yarkoni et al., 2011; Rubin et al., 2017). This “decoding” approach with reverse inference raises hypotheses about the putative cognitive processes underpinning intentional behaviour, where different cognitive processes are associated with specific networks of the human brain. We then reviewed results from these meta-analyses in the context of current cognitive models of intentional choice.

## 2. Materials and Methods

### 2.1 Study selection and inclusion criteria

We defined intentional choices as experimental paradigms involving self-initiated, voluntary selections of an action from two or more alternatives (Zhang et al., 2012). The experimental procedure would need to instruct participants that there are no correct or incorrect choices, and they are free to choose any option among available alternatives. This type of intentional choices differs from conventional goal-directed or externally cued behaviour, in which a correct or instructed response could be defined or identified. We focused on existing studies investigating the *“what”* (which action to choose) or “*whether*” (whether or not to execute an action) component of intentional behaviour (Brass and Haggard, 2008). Studies focusing on the “*when*” component (i.e., when to execute, as in Libet et al., 1983) is not considered here, because of the low temporal resolution of haemodynamic and metabolic responses.

We conducted a systematic literature search in accordance with the PRISMA guidelines (Moher et al., 2015) to identify brain imaging studies of intentional choice. The literature search was performed on both PubMed and PubMed Central (PMC) databases, because the two databases may contain different publications. The PubMed database was searched with specified keywords as following: (“volitional decision” OR “volitional choice” OR “voluntary decision” OR “intended decision” OR “intentional decision” OR “voluntary choice” OR “intended choice” OR “intentional choice” OR “free decision” OR “free choice” OR “volitional action” OR “voluntary action” OR “intended action” OR “intentional action” OR “free action” OR “volitional selection” OR “voluntary selection” OR “intended selection” OR “intentional selection” OR “free selection”) AND (“fMRI” OR “functional Magnetic Resonance Imaging” OR “BOLD” OR “Blood Oxygen Level-Dependent” OR “Positron Emission Tomography”). For the PMC database, the same keywords were employed in the interrogation and a filter on the search field was set to “Body - Key Terms” to constrain search in a more concrete range. The search results from PubMed and PMC databases were combined with duplicated records removed, resulting in 332 publications as of October 2020.

We then inspected every publication from the literature search. The further inclusion criteria for our meta-analysis were applied as follows:

1. Studies reported first handed data that comes from experiments rather than reviews or meta-analysis. 291 of the 332 publications met this criterion.
2. Studies included results from healthy adult human participants. 239 of the remaining 291 publications met this criterion.
3. Studies employed an intentional choice paradigm(s) and reported a fMRI/PET contrast of intentional choice vs. specified response conditions. Here, in the specified response condition, participants responded with the same set of possible actions as in the intentional choice condition, but the identity of which action to respond (or whether to respond) was determined by the experimenter. 40 of the remaining 239 publications met this criterion.
4. Studies reported whole-brain analysis with MNI or Talairach coordinates of the cluster peaks. 37 of the remaining 40 publications met this criterion.
5. If more than one appropriate contrast with the same group of subjects were reported in a single study, only one contrast was included in the meta-analysis.

### 2.2 Activation Likelihood Estimation (ALE) meta-analysis of intentional decision

After the screening, the 37 fMRI/PET studies met the selection criteria, which included 39 independent experiments for meta-analysis. 27 studies recruited only right-handed participants, 1 study recruited thirteen right-handed and one left-handed participants, and the other 9 studies did not specify participants’ handedness. These studies contained a total of 685 participants and reported 344 peak foci of increased fMRI/PET responses to the intentional choice vs. specified response contrast. Less than 3% foci (10 out of 344) were out of the brain mask in the GingerALE toolbox (Turkeltaub et al., 2002; Eickhoff et al., 2012), which was within the normal range due to spatial smoothing and potential registration errors (Eickhoff et al., 2012). Therefore, all foci were included in the study to maximize the usage of the original dataset. For activation foci reported in the Talairach space, we converted them to MNI coordinates using the Lancaster algorithm (Lancaster et al., 2007).

The coordinates-based activation likelihood estimation (ALE) meta-analysis was conducted over all the 38 experiments using the Ginger-ALE toolbox (www.brainmap.org, version 3.0.2) (Turkeltaub et al., 2002; Eickhoff et al., 2012). This analysis aimed to determine, across independent experiments, significant spatial convergence of fMRI/PET activation probabilities for the intentional choice vs. specified response contrast, under the null hypothesis that the activation foci are distributed randomly throughout the brain. First, for each experiment, the activation probabilities of all foci reported were modelled as 3D Gaussian probability distributions with their full-width half-maximum (FWHM) estimated from the between-subject variance of the experiment (Eickhoff et al., 2009). Second, an ALE activation map was then calculated by combining all experimental-level activation maps, yielding a voxel-wise ALE score to quantify the convergence of results across experiments at each voxel location. Third, an analytical approach was used to determine the null distribution of voxel-wise ALE scores. A non-parametric *p*-value map of ALE scores was then generated under the null distribution (Eickhoff et al., 2012). Finally, the *p*-value map was thresholded at *p<*0.001 and corrected for multiple comparisons across voxels using a cluster-level family-wise error (FWE) correction from 5,000 permutations (*p*<0.01, cluster-corrected).

### 2.3 Paradigm-specific meta-analysis

We categorized the 39 experiments into four intentional choice paradigms based on their experimental designs and procedures (Figure 1). The first category of paradigm is referred to as reactional intention (RI), in which participants voluntarily choose cues that associate to specific motor actions. We considered this category as the simplest form of intentional choice because a cue in the RI paradigm is directly linked to a target action. The second category is referred to as perceptual intention (PI), in which participants voluntarily choose between perceptually distinct targets (e.g., icons or pictures). Compared to the RI paradigm, the PI paradigm involves an additional matching process: an option is associated with a perceptual target, and the target is then associated with a specific motor action. The third category is referred to as inhibitory intention (II), in which at least one option is not to act (i.e., withholding responses). A cue in the II paradigm is associated directly with a specific action or the inhibition of action. The final category is referred to as cognitive intention (CI). The free choice condition in CI paradigm requires the participants to choose between options that require higher-order cognitive processes such as doing arithmetic or generating words.

For studies employed each of the four paradigm categories, we performed the same ALE meta-analysis to identify the spatial convergence of fMRI/PET activation for the intentional choice vs. specified response contrast. The same procedure to correct for multiple comparisons was applied as in the meta-analysis across all studies (see section 2.2).

Based on the thresholded ALE maps from individual paradigms, we then conducted further conjunction and contrast meta-analyses between the RI and PI paradigms as well as the RI and II paradigms, using the “contrast study” function implemented in GingerALE. This allowed us to localize voxels commonly (i.e., conjunction) or differentially (i.e., contrast) activated across intentional choice paradigms. The conjunction images were created using the voxel-wise minimum value of the input ALE images, while the contrast images are created by directly subtracting one input image from the other. Because the contrast images unavoidably contained only a subset of studies, similar to previous meta-analyses (Zapparoli et al., 2017), a more lenient threshold (cluster threshold 200 mm^3^, uncorrected voxel-level threshold *p*<0.01, permutation tests with 5,000 iterations) was applied to the contrast analyses between paradigm categories to avoid type II errors (Lieberman and Cunningham, 2009). No conjunction or contrast meta-analysis was conducted on experiments using the CI paradigm due to the limited number of studies available in that category.

### 2.4 Meta-analytic decoding of intentional decision

ALE activation maps indicate brain regions of consistent fMRI/PET activations between studies. We then used NeuroSynth (Yarkoni et al., 2011) to further perform a “reverse-inference” type of meta-analysis. That is, we meta-analytically decoded which cognitive functions or processes are likely to give rise to the consistent brain activations observed in ALE activation maps. As highlighted previously, one should interpret results from reverse inference with caution (Poldrack, 2011). Most functional brain imaging results are correlational. The involvement of a brain region in a certain cognitive function does not directly support the notion that the region is exclusively associated with the cognitive process. Nevertheless, meta-analytic decoding against large, unbiased imaging databases did provide useful information about the engagement of cognitive processes (Poldrack, 2006). In the current study, we consider our meta-analytic decoding analysis to be contributory rather than confirmatory, which offers insights for future studies of intentional decision.

We considered a set of 100 cognitive topics that were previously generated from over 11,000 brain imaging studies. The 100 topics were extracted by fitting a generative statistical model of sematic topics (Blei et al., 2003) to the abstracts of over 11,000 brain imaging articles in the NeuroSynth database (for details see Poldrack et al., 2012). We ignored the topics related to general methods (e.g., fMRI) and focus only on the topics related to cognitive processes. For each cognitive topic, a whole-brain association-test map (also referred to as the reverse inference map) was generated from all the articles in the database. The value at each voxel of the association-test map quantifies the extent to which studies loaded highly on the current topic reported more consistent activation at this location than all the other studies (Yarkoni et al., 2011).

We estimated the similarity between each unthresholded ALE activation map with respect to the association-test maps of the 100 cognitive topics by calculating their Pearson correlations across voxels. The resulting correlation coefficients were rank-ordered to identify the cognitive topics that are most likely to be present during intentional decision and its specific paradigms.

## 3. Results

### 3.1 Meta-analysis of intentional decision

Thirty-five brain imaging studies were identified from our symmetric literature search, which included 38 independent experiments of intentional decision. The number of participants, experimental paradigms and other details were summarized in Table 1.

**Table 1.** List of intentional decision studies that meet the inclusion criteria.

| No. | Study | Number of subjects | Imaging modality | Experiment Paradigm |
| --- | --- | --- | --- | --- |
| 1 | (Beudel and De Jong, 2009) | 16 | fMRI | RI: Press button with 2nd to 5th finger of one hand |
| 2 | (Deiber et al., 1991) | 8 | PET | RI: Push the joystick |
| 3 | (Deiber et al., 1996) | 13 | PET | RI: Move finger in response to visual cue |
| 4 | (François-Brosseau et al., 2009) | 14 | fMRI | RI: Press button with 2nd to 5th finger of one hand<sup>▲</sup> |
| 5 | (Frith et al., 1991) | 6 | PET | RI: Move finger in response to touch |
| 6 | (Gerardin et al., 2004) | 9 | fMRI | RI: Press button with left or right thumb<sup>▲</sup> |
| 7 | (Hoffstaedter et al., 2012) | 35 | fMRI | RI: Press button with right or left hand |
| 8 | (Hyder et al., 1997) | 9 | fMRI | RI: Move finger in response to touch |
| 9 | (Krieghoff et al., 2009) | 16 | fMRI | RI: Press left or right button with the index finger of one hand |
| 10 | (Mueller et al., 2007) | 16 | fMRI | RI: Press right or left button with the index finger of one hand |
| 11 | (Rae et al., 2014) | 17 | fMRI | RI: Press button with 2nd to 5th finger of one hand |
| 12 | (Rowe et al., 2010) | 20 | fMRI | RI: Press button with 2nd to 5th finger of one hand |
| 13 | (Schouppe et al., 2014) | 22 | fMRI | RI: Modified flanker paradigm |
| 14 | (Van Eimeren et al., 2006) | 12 | fMRI | RI: Press button with 2nd & 3rd finger of both hands |
| 15 | (Van Oostende et al., 1997) | 2 | fMRI | RI: Press button with 2nd to 5th finger of one hand |
| 16 | (Walton et al., 2004) | 9 | fMRI | RI: Press button with 2nd to 4th finger of one hand |
| 17 | (Walton et al., 2004) | 9 | fMRI | RI: Press button with 2nd to 4th finger of one hand |
| 18 | (Zapparoli et al., 2017) | 32 | fMRI | RI: Press button with right or left finger |
| 19 | (Bode et al., 2013) | 15 | fMRI | PI: Choose picture by button pressing |
| 20 | (Filevich et al., 2013) | 23 | fMRI | PI: Choose number with mouse cursor |
| 21 | (Forstmann et al., 2006) | 22 | fMRI | PI: Choose target(s) by button pressing |
| 22 | (Lau et al., 2004b) | 12 | fMRI | PI: Choose target(s) by button pressing |
| 23 | (Orr and Banich, 2014) | 28 | fMRI | PI: Choose task by button pressing |
| 24 | (Rens et al., 2018) | 24 | fMRI | PI: Choose target door by button pressing |
| 25 | (Rowe et al., 2005) | 12 | fMRI | PI: Choose target(s) by button pressing |
| 26 | (Rowe et al., 2008) | 20 | fMRI | PI: Choose target(s) by button pressing |
| 27 | (Thimm et al., 2012) | 28 | fMRI | PI: Choose picture by button pressing |
| 28 | (Dall'Acqua et al., 2018) | 24 | fMRI | II: Adapted go/no-go paradigm* |
| 29 | (Karch et al., 2010b) | 8 | fMRI | II: Adapted go/no-go paradigm* |
| 30 | (Karch et al., 2010a) | 15 | fMRI | II: Adapted go/no-go paradigm* |
| 31 | (Karch et al., 2009) | 14 | fMRI | II: Adapted go/no-go paradigm* |
| 32 | (Lynn et al., 2016) | 21 | fMRI | II: Pain stop or endurance by button pressing or not |
| 33 | (Omata et al., 2019) | 26 | fMRI | II: Whether to stop the continuous finger-tapping |
| 34 | (Schel et al., 2014) | 24 | fMRI | II: Adapted go/no-go paradigm* |
| 35 | (Frith et al., 1991) | 6 | PET | CI: Generate word or repeat word |
| 36 | (Jarvstad and Gilchrist, 2019) | 23 | fMRI | CI: saccadic selection |
| 37 | (Ort et al., 2019) | 22 | fMRI | CI: Redirect attention to target(s) without actual movement |
| 38 | (Taylor et al., 2008) | 18 | fMRI | CI: Redirect attention to target(s) without actual movement |
| 39 | (Wisniewski et al., 2016) | 35 | fMRI | CI: mathematical calculation (subtract or add) |
| Total | / | 685 | / | / |

\* All are adapted go/no-go task which has an intentional trial in addition to the tradition go/no-go trials. In the intentional trial, participants are free to choose whether to press the button.
^ The study reported the contrast of intentional decision and specified response separately for left and right hands. Only the results from the dominant hand (right hand) were included in the meta-analysis.

Across all the 39 studies, a Ginger-ALE meta-analysis on the contrast between free choice and specified response yielded greater BOLD-fMRI/PET activations related to intentional behaviour in a frontoparietal network (Figure 2). The analysis identified 17 peaks in 6 clusters, including bilateral pre-supplementary motor area (pre-SMA), bilateral anterior cingulate cortex (ACC), bilateral dorsal lateral prefrontal cortex (dlPFC), bilateral supramarginal gyrus (IPL) and left Insula (Table 2, *p*<0.01, cluster-level corrected).

**Table 2.**
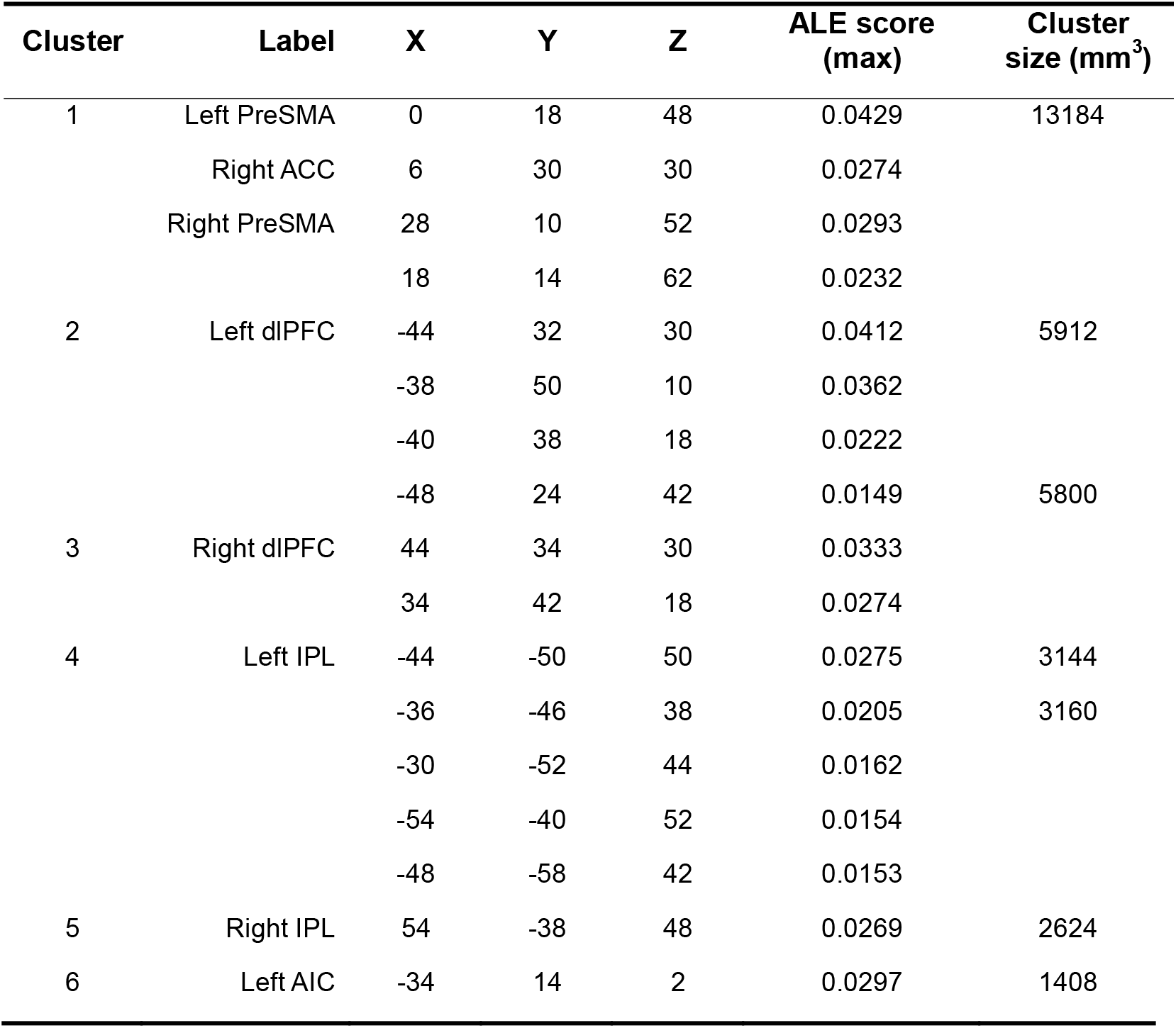
Meta-analysis results of intentional decision (“free choice” > “specified response”) across all studies. Peak coordinates of clusters were reported in the MNI space (mm).

**Figure 2.**
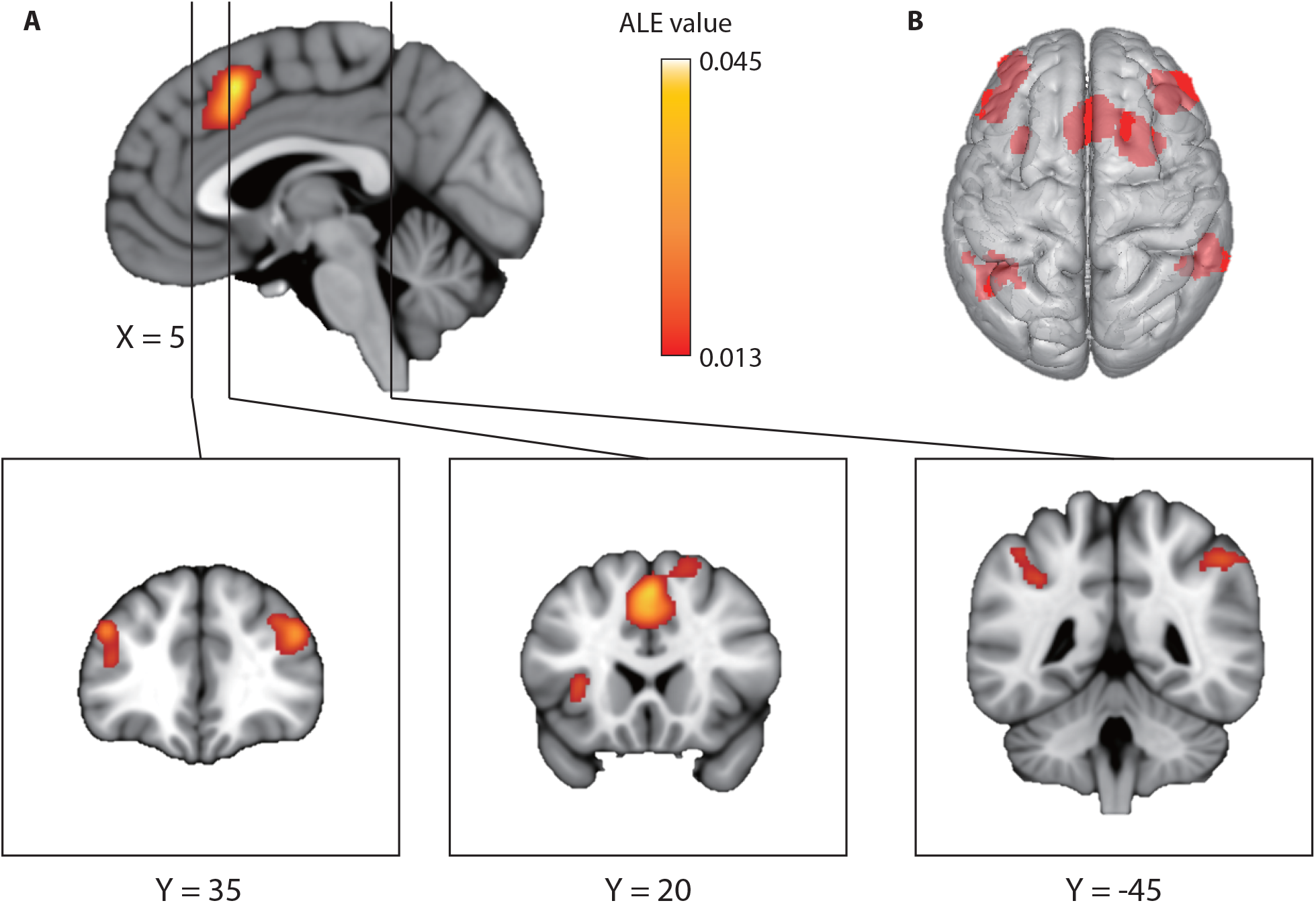
ALE meta-analyses of all free-choice studies showing significant activation clusters related to intentional decision (p<0.01, cluster-level FWE corrected from 5,000 permutations). (A)ALE value map. (B) 3D render of all the clusters. Table 2 lists the peak coordinate of each cluster.

### 3.2 Meta-analysis of contrasts between intentional decision paradigms

To investigate whether different types of intentional behaviour relate to selective brain responses, we assigned intentional decision studies into four categories (Figure 1), depending on their characteristics of experimental paradigms: reactive intention (RI), perceptual intention (PI), inhibitory intention (II) and other higher cognitive intention (CI).

Separate meta-analyses on individual intentional decision paradigms revealed overlapping and distinct clusters with increased activity in response to intentional behaviour (Table 3). Free choices in the RI paradigm were consistently associated with greater activations in four clusters, including bilateral pre-SMA and ACC, bilateral IPL and left dlPFC. For the PI paradigm, the analysis revealed 5 clusters with 6 peak foci located in bilateral dlPFC, bilateral precuneus and left pre-SMA. For the II paradigm, there were 4 clusters with 6 peak foci located in bilateral IPL, right dlPFC and right premotor area. No significant results were observed in the meta-analysis of the CI paradigm, possibly due to the limited number of studies in that category.

**Table 3.** Meta-analysis results of individual paradigms of intentional decision. Peak coordinates of clusters were reported in the MNI space (mm).

| Cluster | Label | X | Y | Z | ALE score (max) | Cluster size (mm<sup>3</sup>) |
| --- | --- | --- | --- | --- | --- | --- |
| Reactive Intention |  |  |  |  |  |  |
| 1 | Bilateral PreSMA | 0 | 18 | 48 | 0.0212 | 4952 |
|  | Left ACC | -6 | 20 | 38 | 0.0203 |  |
|  | Right ACC | 6 | 24 | 42 | 0.0183 |  |
|  |  | 4 | 30 | 28 | 0.0136 |  |
| 2 | Left dlPFC | -44 | 32 | 24 | 0.0193 | 2568 |
|  |  | -36 | 46 | 14 | 0.0170 |  |
| 3 | Right IPL | 54 | -34 | 50 | 0.0183 | 1176 |
|  |  | 56 | -40 | 38 | 0.0100 |  |
| 4 | Left IPL | -38 | -44 | 38 | 0.0198 | 2032 |
|  |  | -50 | -40 | 48 | 0.0135 |  |
| Perceptual Intention |  |  |  |  |  |  |
| 1 | Right dlPFC | 44 | 34 | 32 | 0.0156 | 1512 |
| 2 | Left PreSMA | -4 | 16 | 48 | 0.0191 | 1056 |
| 3 | Left dlPFC | -44 | 32 | 30 | 0.0143 | 968 |
|  |  | -42 | 24 | 32 | 0.0136 |  |
| 4 | Right IPL | 34 | -62 | 42 | 0.0169 | 960 |
| 5 | Left IPL | -8 | -72 | 54 | 0.0147 | 864 |
| Inhibitory Intention |  |  |  |  |  |  |
| 1 | Right IPL | 54 | -42 | 46 | 0.0168 | 1544 |
| 2 | Left IPL | -44 | -48 | 50 | 0.0193 | 1640 |
|  |  | -54 | -42 | 50 | 0.0090 |  |
| 3 | Right Premotor | 28 | 12 | 56 | 0.0143 | 1264 |
|  |  | 18 | 16 | 64 | 0.0130 |  |
| 4 | Right dlPFC | 44 | 40 | 22 | 0.0138 | 712 |

To quantify the distinction and similarity in brain response to different types of intentional behaviour, we conducted further contrast and conjunction meta-analyses, comparing both the PI paradigm (involving perceptual processing) and the II paradigm (involving inhibitory processing) with the most elementary paradigm (the RI paradigm). The contrast meta-analysis showed that the right angular area is more likely activated in the PI than the RI paradigm, and bilateral IPL is more likely to activate in the II than the RI paradigm (Figure 3A, Table 4). The conjunction meta-analysis showed that bilateral Pre-SMA/ACC complex and the left dlPFC are commonly activated in intentional behaviour across studies of PI and RI paradigms, and activations in the bilateral IPL are commonly observed in both II and RI paradigms (Figure 3B, Table 4).

**Table 4.** Contrast and conjunction meta-analyses between different free-choice paradigms. Peak coordinates of clusters were reported in the MNI space (mm).

| Cluster | Label | X | Y | Z | ALE score | Cluster size (mm<sup>3</sup>) |
| --- | --- | --- | --- | --- | --- | --- |
| **Perceptual Intention > Reactive Intention** |  |  |  |  |  |  |
| 1 | Right Angular gyrus | 38 | -66 | 44 | 0.0125 | 312 |
|  |  | 36.7 | -64.7 | 39.3 | 0.0113 |  |
|  |  | 37.5 | -60 | 42 | 0.0094 |  |
| **Inhibitory Intention > Reactive Intention** |  |  |  |  |  |  |
| 1 | Right IPL | -40.8 | -50.5 | 50.2 | 0.0137 | 448 |
| 2 | Left IPL | 51.2 | -46 | 45.2 | 0.0179 | 1192 |
| **Perceptual Intention ∩ Reactive Intention** |  |  |  |  |  |  |
| 1 | PreSMA/ACC | -2 | 18 | 48 | 0.0179 | 744 |
| 2 | Left dlPFC | -44 | 32 | 30 | 0.0125 | 232 |
| **Inhibitory Intention ∩ Reactive Intention** |  |  |  |  |  |  |
| 1 | Right IPL | 54 | -38 | 46 | 0.0128 | 360 |
| 2 | Left IPL | -54 | -42 | 52 | 0.0090 | 40 |

**Figure 3.**
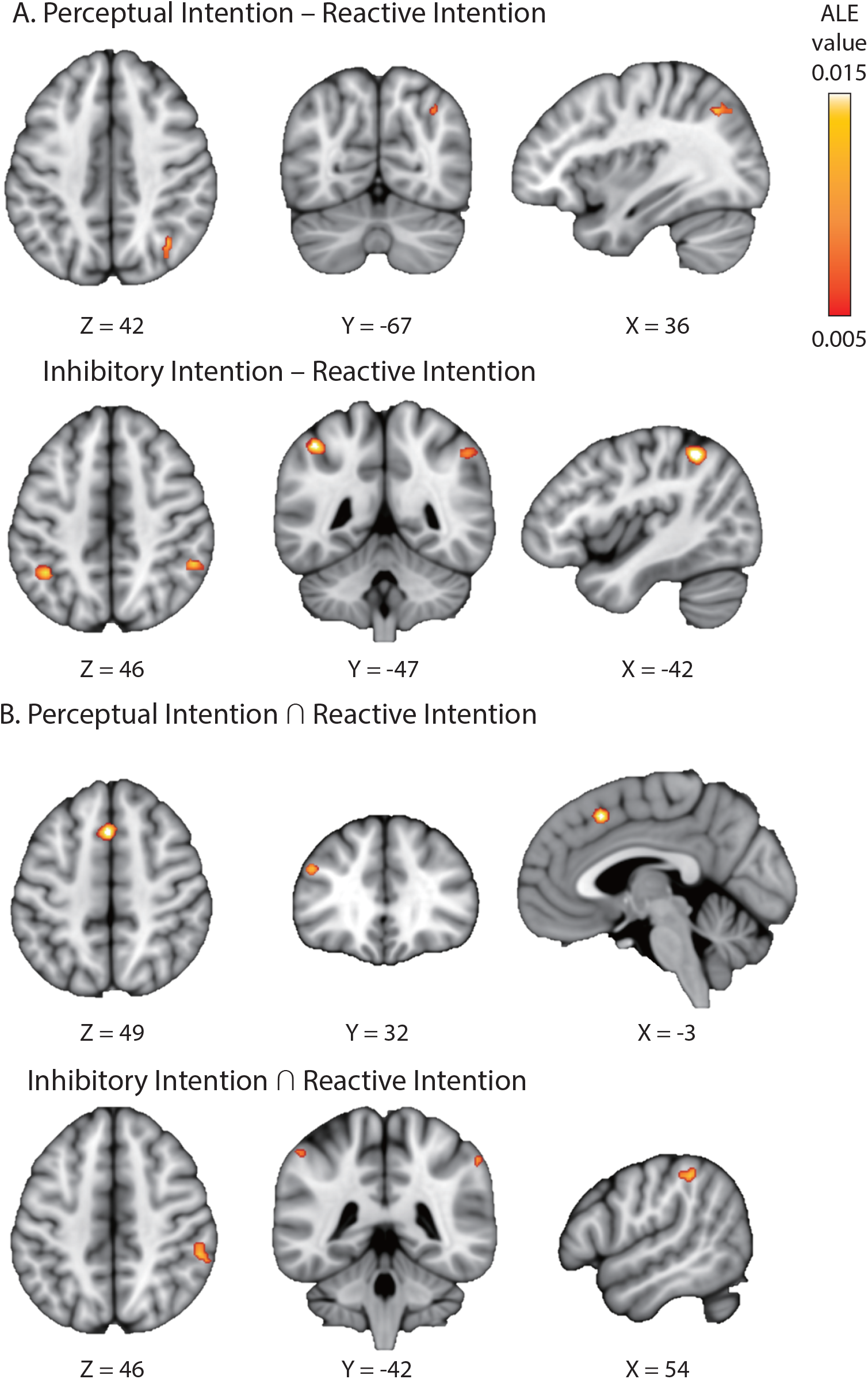
Contrast and conjunction meta-analyses (A) ALE contrast meta-analyses between PI vs. RI (top) and II vs. RI (bottom) paradigms. (B) ALE conjunction meta-analyses between PI and RI paradigms (top) as well as II and RI paradigms (bottom). Table 4 lists the peak coordinate of each cluster.

### 3.3 Meta-analytic decoding of intentional decision

To probe cognitive processes underlying intentional decision, we assessed the spatial similarity (i.e., Pearson correlation across voxels) between the ALE activation maps from our meta-analysis and 100 association-test maps. Each of the association-test maps represents brain response selective to one of 100 psychological topics, generated from meta-analyses of >11,000 independent studies (Yarkoni et al., 2011). Therefore, a high correlation coefficient to an association-test map would imply the potential involvement of the corresponding cognitive process. The primary interest here is the relative ranking of the topics based on the similarity of their association-test maps to our results, not to perform null hypothesis significance testing on each correlation.

This reverse inference showed that the frontoparietal network identified in the meta-analysis of intentional decision across all studies (Figure 4) were strongly associated with several psychological topics. The top three are working memory (*R* = 0.434), task rules (*R* = 0.394) and conflict (*R* = 0.367) (Figure 4, and see Supplementary Table 1 for the full results).

**Figure 4.**
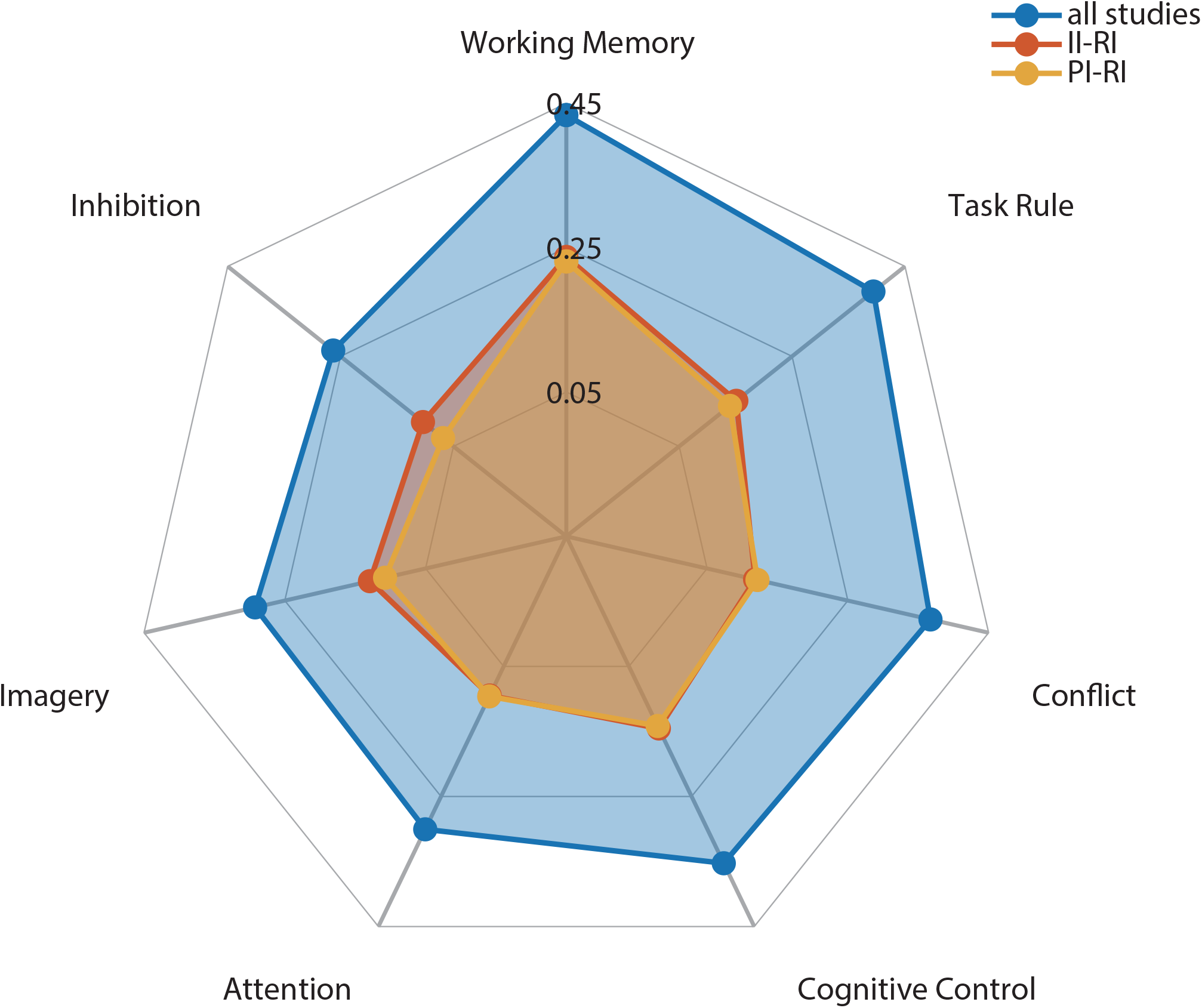
Meta-analytic decoding of intentional decision. Correlation coefficients between different cognitive topics’ association maps and ALE maps of intentional decision were calculated. The correlation values for the top 7 topics were illustrated in a polar plot. Terms used to generate those topic-based association maps were listed in Supplementary Table 1.

The ALE-activation map from the meta-analytical contrast between PI and RI paradigms (Figure 3A) was associated with topics of working memory (*R* = 0.231), cognitive control (*R* = 0.142) and switching rules (*R* = 0.140). The meta-analytical contrast between II and RI paradigms was associated with topics of working memory (*R* = 0.237), switching rules (*R* = 0.151), conflicts (*R* = 0.145) and response inhibition (*R* =0.104).

## 4. Discussion

This study confirms, at the meta-analytic level, a consistent pattern of brain activation associated with intentional decisions for action, perception and cognition. By contrasting between different free-choice paradigms, we also identified brain areas whose activities are dependent on specific categories of intentional behaviour. The meta-analytic decoding analysis suggested putative cognitive processes underlying intentional decisions. Our results provide insight into the cognitive roles of brain networks that mediate intentional behaviour in humans, which we discuss below together with their potential computational processes.

### 4.1 Functional localization of intentional decision in the brain

The ALE meta-analysis showed increased activities in the medial prefrontal cortex (pre-SMA and caudal ACC), the lateral frontoparietal cortices (DLPFC and IPL) and the anterior insula cortex (AIC) during voluntary behaviour originated from intentional decisions, in contrast of the same behavioural response prescribed exogenously (Figure 2). This result is in agreement with previous analyses that applied a similar method to smaller study samples (Rae et al., 2014; Zapparoli et al., 2017).

The brain areas involved in intentional decision overlap closely with the multiple demand network (Duncan and Owen, 2000; Duncan, 2010), a “task-positive” co-activation pattern associated with diverse cognitive demands (Fox et al., 2005; Dosenbach et al., 2006). A closer inspection of the literature indicates that subcomponents of this network may serve different cognitive roles during intentional decisions, which is also supported by our meta-analytic decoding results (Figure 4).

A large body of evidence indicates the central role of ACC in conflict monitoring (Botvinick et al., 2004). Conflicts in information processing arise from the presence of response competition. Greater ACC activation is consistently observed when (1) one or more prepotent responses need to be overridden, such as in the Stroop task (MacLeod and MacDonald, 2000; Barch, 2001) and the flanker task (Botvinick et al., 1999; Bunge et al., 2002), or (2) a voluntary choice is needed among multiple underdetermined options, like in all the free-choice paradigms discussed here. Although the existing literature of conflict monitoring is largely focused on the ACC, the adjacent pre-SMA is also sensitive to the presence of conflict, in particular the conflict in response selection, as lesions in this region lead to deficits in exerting voluntary control over immediate actions (Nachev et al., 2007). According to the conflict monitoring theory, as multiple options become available in the free-choice paradigm, increased ACC and pre-SMA activities may encode conflicts as an index of the need for greater cognitive demand, which in turn trigger voluntary choices to reduce or resolve the conflict (Botvinick et al., 2004; Botvinick, 2007). A direct prediction of this proposition is that the activity in the medial prefrontal cortex should increase proportionally, at least to some extent, to the number of available options in the free-choice paradigm, which has been validated in previous studies (e.g., Forstmann et al., 2006).

Beyond the medial prefrontal cortex, the frontoparietal network on the lateral brain surface has a distinct functional connectivity pattern relating to cognitive control (Corbetta and Shulman, 2002) and executive task performance (Seeley et al., 2007). Two functions of this network are essential to intentional behaviour. First, intentional decisions in the free-choice paradigm are, by definition, rendered endogenously. Nevertheless, the brain may still establish a “task set” that incorporates transient and arbitrary rules in addition to relevant exogenous information, such as associations of stimuli and imagined outcomes as well as available options (Sakai, 2008). Both single-unit recording in non-human primates (Quintana and Fuster, 1999; Asaad et al., 2000; Wallis et al., 2001) and brain imaging in humans (Bunge et al., 2002; Sakai and Passingham, 2003) have identified neural representations of various task sets in the frontoparietal network. The encoding of a task set can be actively maintained in this network until its execution (Zhang et al., 2013), thereby facilitating the intentional decision process to unfold in time. Second, intentional behaviour is commonly accompanied by the subjective experience of volitional control (Haggard, 2008), which requires internal models that matches the consequences of the response against its initial intention (Wolpert et al., 1995). It has been proposed that the parietal cortex hosts such internal models (Desmurget and Grafton, 2000), as patients with parietal lesions exhibited altered behavioural and electrophysiological signatures of their intention to act (Sirigu et al., 2004).

Our meta-analysis across all free-choice experiments showed the consistent involvement of the AIC during intentional decision, in spite of the lack of significant insula activity in some studies (e.g., Van Eimeren et al., 2006). This supports an earlier account that the AIC is a key component of the integrated brain network involved in intentional behaviour (Brass and Haggard, 2010). Anatomically, the AIC connects directly to the ACC (Augustine, 1996; Moisset et al., 2010). Functionally, robust coactivation in the AIC and ACC was observed across multiple cognitive domains (Medford and Critchley, 2010) as well as in resting-state (Chang et al., 2013), and both regions are a part of the salience network (Chen et al., 2016). It may therefore be tempting to ascribe the AIC activity to conflict processing during intentional decision, similar to that of ACC. An alternative proposal originated from the AIC’s unique function in signalling introspective awareness (Craig, 2009) or subjective salience (Menon and Uddin, 2010) of cognitive (Preuschoff et al., 2008), homeostatic (Craig et al., 2000; Farrer and Frith, 2002) and emotional (Jabbi et al., 2007) information, which is not shared with the ACC. According to this theory, AIC activity reflects the affective consequences of intentional decisions. In other words, the AIC may not directly associate with the formation of current intention; instead, it evaluates the outcome of the intentional act with respect to an internal model of one’s long term goal (see Brass and Haggard, 2010 for a detailed review).

### 4.2 Computational processes of intentional decision

With the identification of the consistent brain network for intentional decision-making, a new question arises: what is the computational process underlying intentional decision? Converging findings from behavioural modelling (Ratcliff, 2006), single-unit recoding (Kim and Shadlen, 1999; Shadlen and Newsome, 2001; Mazurek et al., 2003) and imaging (Heekeren et al., 2004; Ploran et al., 2007; Ho et al., 2009) experiments suggest that, when making choices based on external stimuli, an accumulation-to-threshold mechanism governs the decision-making process (Smith and Ratcliff, 2004; Gold and Shadlen, 2007; Heekeren et al., 2008): the evidence supporting one or multiple options are accumulated over time, and a choice is made when the accumulated evidence reached a decision threshold. For perceptual decisions with noisy sensory stimuli, this accumulation process reduces the momentary noise in information-processing and in turn results in more accurate decisions (Bogacz et al., 2006, 2007; Zhang and Bogacz, 2010).

For intentional decisions, it has been shown that a computational model implementing the accumulation-to-threshold mechanism can well describe the behavioural performance (i.e., response time distributions and choice probabilities) of both RI (Zhang et al., 2012) and PI paradigms (Zajkowski et al., 2020). Furthermore, the accumulated evidence predicted by the model is associated with the BOLD response in the ACC and pre-SMA on a trial-by-trial basis (Zhang et al., 2012). These results raise an intriguing possibility that, during intentional decision, the medial prefrontal cortex implements the accumulation-to-threshold process to integrate over time the transitory intention of choosing different options, until the accumulated intention for one choice reaches a decision threshold.

This hypothesis is supported by several electrophysiological studies, which characterised the accumulation process during intentional behaviour at a high temporal resolution. First, in Libet’s paradigm of voluntary action, the readiness potential measured by scalp EEG precedes participants’ conscious awareness of their voluntary intention (i.e., the “urge to move”, Libet et al., 1983). An accumulator model can be fit to the time latency of participants’ urge to move, and the activity of the accumulator qualitatively reproduces the time course of the readiness potential prior to conscious intention (Schurger et al., 2012). Second, in a free-choice version of Libet’s paradigm, when participants made intentional decisions between responding with their left or right hands, single-neuron activity in the medial prefrontal cortex build up several hundred milliseconds before the onset of conscious intention (Fried et al., 2011). Further, medial prefrontal neurons contralateral to the acting hand exhibited larger activity than ipsilateral neurons (Fried et al., 2011). Therefore, the medial prefrontal cortex may host accumulated intentions of multiple responses as well as their mutual competition, from which voluntary acts are rendered via the accumulation-to-threshold mechanism.

The putative role of the medial prefrontal cortex in intention accumulation is not inconsistent with this region’s function of conflict monitoring discussed above, because more free options would be associated with larger accumulated intention across alternatives as well as higher conflict. In this regard, intention accumulation can be interpreted as a computational implementation of detecting and resolving conflicts among underdetermined options. Therefore, we consider the accumulation process as a parsimonious computational framework for intentional behaviour outlined by the conceptual *what*-*when*-*whether* model (Brass and Haggard, 2008), because accumulator models can explain quantitatively both “*what”* (i.e., choice probabilities) and *“when”* (i.e., response time distributions) components. Interestingly, accumulator models can also be fitted to behavioural performance in externally-triggered stopping tasks (Gomez et al., 2007; Zhang et al., 2016). Future research should investigate if accumulator models can incorporate the “*whether*” component, or voluntary stopping in the II paradigm.

### 4.3 Paradigm-specific activations during intentional decision

By categorizing free-choice studies into different types, we identified brain regions associated with consistent and specific activations between sub-categories of intentional decision (Figure 3). The conjunction meta-analysis of the RI and PI paradigms showed that the pre-SMA/ACC and DLPFC are associated with both types of intentional decision. This is expected, as the RI and PI paradigms have a similar task structure, involving rapid voluntary choices among multiple action plans. On the other hand, the contrast meta-analysis between the two suggested that, compared with the simple RI paradigm, intentional behaviour in the PI paradigm more likely involve the right angular gyrus (AG). The AG plays an important role in reorienting and shifting of attention (Seghier, 2013), as well as in updating attention allocation with task-related information (Ciaramelli et al., 2008; Taylor et al., 2011). Therefore, the increased recruitment of AG in the PI paradigms could be due to the additional demand of attention re-allocation for matching between perceptual targets and their respondent actions (Figure 1B).

The II paradigm includes a unique option of not to act or intentionally inhibit one’s action (Figure 1C). The conjunction meta-analysis of the II and RI paradigms showed that the bilateral supramarginal gyrus in the IPL was associated with both types of intentional decisions, and the contrast meta-analysis showed that the supramarginal gyrus was more likely to be activated in the II than that in the RI paradigm. In both RI and II paradigms, participants need to reprogram their response model according to available options in each trial, which fits the critical role of the supramarginal gyrus in action reprogramming (Hartwigsen et al., 2012). The same region is also sensitive to the content of actin plans and their similarity (Quandt et al., 2017). It could be argued that options in the II paradigm are more dissimilar (i.e., acting versus stopping) than that in the RI paradigm (i.e., multiple but similar actions), which leads to the additional recruitment of the supramarginal gyrus in the II paradigm.

It is worth noting that the results of conjunction and contrast meta-analyses should be interpreted with caution, because of the limited number of studies available in each category. Furthermore, one potential confound of the contrast meta-analysis is that different paradigm categories may vary in their task difficulty, and hence the contrast between categories may not directly support the involvement of distinct cognitive processes. This issue can be examined in future studies that explicitly manipulate both task difficulty and intentional decision paradigms.

### 4.4 Future directions and conclusion

This analysis leaves open some issues for future research on human intentional decision-making. First, our systematic review identified only four studies in the CI category: two studies included options with attention shifts (Taylor et al., 2008; Ort et al., 2019), one with verbal responses (Frith et al., 1991) and the other one with arithmetic rules (Wisniewski et al., 2016). The small number of CI studies did not yield any significant result in the paradigm-specific meta-analysis, but that may reflect type II error. We recommend future research to explore different types of CI studies and examine the robustness and consistency of existing results across a range of distinct cognitive processes.

Second, our meta-analysis of the II paradigm did not show conventional regions involved in inhibitory control (Swick et al., 2011). We propose that this is due to the fact that our analysis used the contrast of intentional choice vs. specified response, with the former including intentional stopping and the latter including externally triggered stopping - this contrast may therefore not detect differential response inhibition. Indeed, the BOLD response in the AIC was higher during intentional stopping than intentional action execution (Brass and Haggard, 2007), while the inferior frontal gyrus is consistently observed during instructed stopping (Aron et al., 2004). To examine how the brain switches effectively between intentional and instructed stopping in the II paradigm, one need to examine the effective connectivity between these two regions and the medial prefrontal cortex, which is involved in both types of stopping (Kühn and Brass, 2009; Sharp et al., 2010).

Third, the current imaging literature on intentional behaviour indicates that the main focus is to localize associated brain regions or their underlying computational processes. Less is known about why a participant would intentionally choose one option over others in a trial. The answer to this question is important because the sequence of intentional decisions over trials are not completely random (Zhang and Rowe, 2015) but dependent on executive control of working memory (Baddeley et al., 1998), the context of a given choice in a sequence (Rowe et al., 2010), and other sources of response biases (Zajkowski et al., 2020). We suggest that the free-choice paradigm provides an ideal testbed for future research to investigate the interplay between the intention accumulation process governing a single trial and modulatory effects that operate at a longer time span.

In conclusion, our meta-analysis identifies a brain network consistently activated when humans have the freedom to make intentional choices among multiple options. Some components of this network are recruited specifically in subcategories of the free-choice paradigm. Multiple cognitive and computational processes are involved in intentional decision, which collectively serve essential roles in shaping and maintaining volitional control.

## Acknowledgements

This study was supported by a European Research Council starting grant to JZ (716321). RS was supported by a PhD studentship from China Scholarship Council. JBR is supported by the Medical Research Council (SUAG/051 G101400) and James S McDonnell Foundation (21st Century Science Initiative, “Understanding Human Cognition”).

## Footnotes

1 In the literature, several terms have been used to refer to the free-choice paradigm, such as “voluntary selection” (Forstmann et al., 2006), “willed action” (Lau et al., 2004b), “internal selection” (Van Oostende et al., 1997), “self-initiated” (Cunnington et al., 2002), and “chosen actions” (Zhang et al., 2012). The current study uses these terms interchangeably.

