## Supplementary Tables for "Functional localization and categorization of intentional decisions in humans: a meta-analysis of brain imaging studies"

### Supplementary Material

Ruoguang Si, James B Rowe, Jiaxiang Zhang

**Supplementary Table 1.** Top 7 cognitive topics and their corresponding correlation coefficients with brain maps from the main (Figure 2) and contrast (e.g., II vs. RI paradigms and PI vs. RI paradigms, Figure 3) meta-analyses. The ranking is based on the meta-analytic decoding result of the main analysis of all studies (Figure 2).

| Topics | All studies | II-RI | PI-RI |
|---|---|---|---|
| Working memory | 0.4337 | 0.2370 | 0.2309 |
| Task rules | 0.3941 | 0.1512 | 0.1399 |
| Conflicts | 0.3674 | 0.1182 | 0.1216 |
| Cognitive control | 0.3527 | 0.1454 | 0.1416 |
| Attention | 0.3005 | 0.0947 | 0.0961 |
| Imagery | 0.2922 | 0.1290 | 0.1075 |
| Inhibition | 0.2630 | 0.1040 | 0.0684 |

**Supplementary Table 2.** Full list of terms associated with the 7 topics in Supplementary Table 1. The number of studies of each topic was extracted from the Neurosynth database (Yarkoni et al., 2011, https://neurosynth.org/analyses/topics/v4-topics-100/).

| Topics | Terms included | Num. Studies |
|---|---|---|
| Working memory | memory, working, task, load, verbal, maintenance, performance, cognitive, activation, information, tasks, term, capacity, probe, manipulation, executive, spatial, phase, encoding, updating, performed, storage, network, span, rehearsal, retention, increased, delay, function, accuracy, functions, demands, vwm, delayed, phonological, loads, demand, performers, sternberg, binding | 798 |
| Task rules | switching, set, rule, task, switch, rules, flexibility, shifting, sets, sorting, trials, shift, shifts, anxiety, switches, card, wcst, anxious, costs, paradigm, single, ef, required, wisconsin, stimulus, trial, worry, repeat, switched, paradigms, execution, depending, component, chunk, cost, gad, lower, updating, types, iu | 230 |
| Conflicts | conflict, interference, control, stroop, incongruent, task, response, congruent, olfactory, resolution, trials, odor, behavioral, attentional, color, cognitive, odors, simon, congruency, word, flanker, effect, monitoring, irrelevant, processing, activated, conflicting, incompatible, neutral, relevant, mechanisms, detection, conflicts, situations, resolve, adjustments, pre, compatibility, compatible, counting | 392 |
| Cognitive control | cognitive, control, performance, task, executive, function, functions, cognition, ability, attention, behavioral, tasks, functioning, test, goal, effort, individuals, behavior, demands, recruitment, abilities, lateral, performed, attentional, neuropsychological, directed, domains, tests, level, individual, relevant, behavioural, evidence, speed, performing, impairment, stroop, deficits, impaired, domain | 1098 |
| Attention | attention, attentional, visual, spatial, search, orienting, target, top, selective, control, location, areas, attended, network, stimulus, irrelevant, cues, distraction, shifts, relevant, feature, modulation, task, cued, mechanisms, color, cueing, focus, bottom, processing, attend, event, endogenous, cue, attending, allocation, directed, resources, modulated, perceptual | 841 |
| Imagery | wm, imagery, mental, imagined, rotation, mi, tasks, visual, visuospatial, motor, areas, spatial, ltm, imagination, imagine, transformation, image, imagining, mentally, images, activated, ability, angle, degrees, physical, strategy, manipulation, visuo, poor, actual, rotations, simulation, kinesthetic, gifted, representational, clock, angles, rehearsal, future, instructed | 355 |
| Inhibition | inhibition, response, control, inhibitory, stop, task, motor, signal, activation, trials, nogo, suppression, responses, successful, inhibit, behavioral, error, inhibited, inhibiting, pre, performance, prepotent, reactive, stopping, ability, sst, monitoring, suppress, correlates, gating, action, rifg, success, proactive, errors, tasks, required, participants, voluntary, behavior | 421 |
